# The small RNA ErsA plays a role in the regulatory network of *Pseudomonas aeruginosa* pathogenicity in airways infection

**DOI:** 10.1101/2020.06.22.164558

**Authors:** Silvia Ferrara, Alice Rossi, Serena Ranucci, Ida De Fino, Alessandra Bragonzi, Cristina Cigana, Giovanni Bertoni

## Abstract

Bacterial small RNAs play a remarkable role in the regulation of functions involved in host-pathogen interaction. ErsA is a small RNA of *Pseudomonas aeruginosa* that contributes to the regulation of bacterial virulence traits such as biofilm formation and motility. Shown to take part in a regulatory circuit under the control of the envelope stress response sigma factor σ^22^, ErsA targets post-transcriptionally the key virulence-associated gene *algC*. Moreover, ErsA contributes to biofilm development and motility through the post-transcriptional modulation of the transcription factor AmrZ. Intending to evaluate the regulatory relevance of ErsA in the pathogenesis of respiratory infections, we analyzed the impact of ErsA-mediated regulation on the virulence potential of *P. aeruginosa* and the stimulation of the inflammatory response during the infection of bronchial epithelial cells and a murine model. Furthermore, we assessed ErsA expression in a collection of *P. aeruginosa* clinical pulmonary isolates and investigated the link of ErsA with acquired antibiotic resistance by generating an *ersA* gene deletion mutant in a multidrug-resistant *P. aeruginosa* strain which has long been adapted in the airways of a cystic fibrosis (CF) patient. Our results show that the ErsA-mediated regulation is relevant for the *P. aeruginosa* pathogenicity during acute infection and contributes to the stimulation of the host inflammatory response. Besides, ErsA could be subjected to selective pressure for *P. aeruginosa* patho-adaptation and acquirement of resistance to antibiotics commonly used in clinical practice during chronic CF infections. Our findings establish the role of ErsA as an important regulatory element in the host-pathogen interaction.

**Author summary:** *Pseudomonas aeruginosa* is one of the most critical multi-drug resistant opportunistic pathogen in humans, able to cause both lethal acute and chronic lung infections. Thorough knowledge of the regulatory mechanisms involved in the establishment and persistence of the airways infections by *P. aeruginosa* remains elusive. Emerging candidates as molecular regulators of pathogenesis in *P. aeruginosa* are small RNAs, which act post-transcriptionally as signal transducers of host cues. Known for being involved in the regulation of biofilm formation and responsive to envelope stress response, we show that the small RNA ErsA can play regulatory roles in acute infection, stimulation of host inflammatory response, mechanisms of acquirement of antibiotic resistance and adaptation during the chronic lung infections of cystic fibrosis patients. Elucidating the complexity of the networks regulating host-pathogen interaction is crucial to identify novel targets for future therapeutic applications.

## Introduction

The bacterium *Pseudomonas aeruginosa* is a common pathogen associated with respiratory tract infections in patients with diverse diseases (1-3). *P. aeruginosa* causes fatal acute lung infections in critically ill individuals who are for instance hospitalized, intubated in an intensive care unit, or immune-compromised (e.g. transplant recipients, patients with burns, cancer, and neutropenia, or infected with HIV). In acute pneumonia, *P. aeruginosa* causes necrosis of the lung epithelium and disseminates into the circulation, resulting in septic shock and multiple organ failure. *P. aeruginosa* is also a major cause of chronic lung infections in individuals with cystic fibrosis (CF), non-CF bronchiectasis, and chronic obstructive pulmonary disease (COPD). It was shown that long-term *P. aeruginosa* persistence in CF airways triggers tissue remodeling that finally leads to lung function decline and ultimately results in respiratory failure.

Biofilm formation is a well-known essential requisite for *P. aeruginosa* during chronic airways infections (4). However, the relevant role of biofilm aggregation of *P. aeruginosa* on the apical surface of polarized epithelial cells at early time points of acute lung infections has also been pinpointed (5-7), challenging the classical notion that acute infections are associated only with the planktonic lifestyle. Indeed, *P. aeruginosa* initiates most acute infections with a transition from planktonic bacteria to host cell-attached aggregates (7). Initial binding of individual sentinel bacteria at the mucosal barrier through two major adhesins, flagella, and retractile type IV pili (8) leads in few minutes to the recruitment of free-swimming bacteria, with the resultant formation of antibiotic-resistant biofilm-like bacterial aggregates of ten to hundreds of bacteria embedded in an exopolysaccharide (EPS) and extracellular DNA (eDNA) matrix and localized in spots on the host cell surface (5, 6). Surface-bound bacterial aggregates, and not individual bacteria, trigger a dramatic remodeling of the apical membrane, namely the formation of protrusions (6). Apical membrane remodeling is linked with localized nuclear translocation of NF-kB underneath aggregates but not beneath single bacteria (9). This indicates the activation of the innate immune response to bacterial aggregates (9). However, aggregate-induced protrusion formation is necessary, but not sufficient, for activation of the innate immune response (9). Indeed, NF-kB activation and the subsequent production of pro-inflammatory cytokines require both pathogen-induced membrane protrusions and the recognition of pathogen-associated molecular patterns (PAMPs) such as flagellin or lipopolysaccharide (LPS) via the cognate Toll-like receptors (TLRs) (10). Once mucosal colonization is established, *P. aeruginosa* delivers a large battery of virulence factors to cause disease, for instance through the type III secretion system (T3SS) that is also required for the bacterial aggregate-mediated induction of membrane protrusions (6). At this stage of acute infection, all virulence factors participate, at different levels, in the cytotoxicity of *P. aeruginosa* that leads to bypassing the epithelial barrier and then to invasion and systemic dissemination (1). Most of the *P. aeruginosa* invasive functions characteristics of acute infection are selected against in CF chronic infection leading to less virulent but more persistent phenotypes (4, 10).

The two pathogenetic processes associated to the progression of *P. aeruginosa* airways infection towards either rapid and acute systemic dissemination or chronic colonization are complex and depend on the coordinate up- or down-regulation of several virulence lifestyle functions that imply both short- and long-term adaptation to host environment (4, 11, 12). For instance, in the pathogenesis of CF chronic infections, *P. aeruginosa* adapted variants can shape the innate immune response favoring their persistence and contribute to the emergence of CF airway hallmarks (13). *P. aeruginosa* adaptive response leading to pathogenesis relies on a wide, intricate, and “prone to remodeling” regulatory network formed both by transcription factors and post-transcriptional regulators including also small RNAs (sRNAs) (14-16). The dynamicity of this regulatory network is frequently observed during the adaptive radiation of *P. aeruginosa* for long-term persistence in the CF lung environment, where bacteria endure various attacks, encompassing oxidative stresses, immune responses, and prolonged antibiotic treatments. To survive these harsh conditions, initial infecting *P. aeruginosa* clones undergo substantial phenotypic changes that may include slow growth, auxotrophy, virulence attenuation, loss of motility, mucoid capsule, biofilm formation, hypermutability, LPS modifications and antibiotic resistance (17). Analysis of several CF clinical isolates showed that adaptive mutations in about 50 genes are mainly responsible for the convergent molecular evolution towards the above-mentioned phenotypes. Of these genes, common mutations occur in around 15 regulatory genes for transcription factors and are supposed to be at the base of remodeling of the infection regulatory network leading to *P. aeruginosa* adaptation to CF lung (17). These regulatory patho-adaptive mutations also involve genes for alternative sigma factors such as PvdS, σ^54^ (RpoN), σ^22^ (AlgT/U), and its repressor MucA (17, 18). *P. aeruginosa* σ^22^ is the functional homolog of *E. coli* σ^E^ (19) that, along with several σ^E^-regulated sRNAs (20, 21), orchestrates the envelope stress response, which in Gram-negative bacteria is critical for maintaining envelope integrity in the host environment and thus to successfully cause infection (20-22).

Generally, sRNAs are key components of the regulatory networks involved in the adaptive response to the stressful conditions that pathogenic bacteria experience during host infection (16, 20-25). Specific protein-RNA and RNA-RNA interactions in the *P. aeruginosa* adaptive regulatory network have been identified for about 16 sRNAs (16). One such *P. aeruginosa* sRNA is ErsA, a 132 nt long transcript that was described for the first time in a work in which 52 novel sRNAs were identified in PAO1 (26) and PA14 (27), two prototype laboratory strains in which ErsA showed to be similarly expressed under laboratory conditions (28). Later, ErsA expression was shown to be strictly dependent on and responsive to envelope stress by σ^22^ (29). Other infection cues such as temperature shifts from environmental to body temperature and reduced oxygen conditions up-regulate ErsA expression (29). Functional studies showed that ErsA contributes to the regulation of virulence traits such as biofilm formation and motility (29, 30). Phenotypically, the knock-out *ersA* mutant strain forms a flat and uniform biofilm and shows enhanced swarming and twitching capability (30). ErsA influences the dynamics of exopolysaccharide production, and the consequent biofilm formation, via negative post-transcriptional regulation of *algC* mRNA (29). The *algC* gene encodes a key point enzyme that coordinates the alginate biosynthetic pathway and the synthesis of several *P*.□*aeruginosa* polysaccharide exoproducts such as Psl, Pel, LPS, and rhamnolipids (31-33). Like ErsA, the expression of *algC* is also dependent on σ^22^ (34, 35), which generates an incoherent feed-forward loop to fine-tune the expression of the AlgC enzyme. Besides, acting as a positive post-transcriptional regulator, ErsA stimulates exopolysaccharide production and biofilm formation also through the post-transcriptional activation of A*mrZ* (30), a transcription factor known to regulate alginate production and motility, and indicated as a molecular switch that triggers biofilm maturation in *P. aeruginosa*. Moreover, ErsA regulatory activity impacts considerably the *P*.□*aeruginosa* transcriptome. More than 160 genes are differentially expressed in RNA-seq experiments comparing the knock-out *ersA* mutant with the PAO1 wild-type. Among these are genes for biofilm formation and motility regulation that also belong to the AmrZ regulon. Furthermore, other differentially expressed genes in the Δ*ersA* mutant are involved in several aspects of *P. aeruginosa*-host interaction, such as denitrification and nitrate metabolism, nitrate transport, type VI and III secretion systems effectors, energy and carbon metabolism, heat-shock proteins and pyocyanin production (30).

Overall, the ErsA ability to respond to host cues and influence the expression of several virulence-associated genes was thought to play a relevant role during host infection. Also, ErsA was implicated in other aspects of *P. aeruginosa* lifestyle linked to infection processes, such as niche establishment/protection in mixed populations, and antibiotic resistance. Indeed, ErsA was suggested to coordinate biofilm maturation dynamics also during mixed-species biofilm growth (36). In the presence of *Staphylococcus aureus*, ErsA is part of that 0.3% of *the P. aeruginosa* genome which becomes differentially expressed. The increase of its transcription levels suggests a role not only in counteracting agents produced by *S. aureus* but also in modulating the state of the exopolymeric matrix for typical biofilm maturation (36). ErsA was also shown to negatively regulate *oprD* mRNA (37), coding for the OprD porin that is the major channel for entry of the carbapenem antibiotics into the periplasm of *P. aeruginosa*. Coherently with these results, strains lacking ErsA were more susceptible to meropenem than the PAO1 wild-type strain (37).

This study aimed to assess the role of ErsA in the regulatory network of *P. aeruginosa* pathogenicity in the infection of the airways. Here we provide evidence that the ErsA-mediated regulation is relevant during acute infection and contributes to the stimulation of the host inflammatory response. Besides, ErsA could also play a regulatory role during chronic infection, in mechanisms of adaptation and acquirement of antibiotic resistance leading to the typical resilient phenotype of *P. aeruginosa* in the CF airways.

## Results

### ErsA contributes to the regulation of bacterial functions involved in cytotoxicity and stimulation of the pro-inflammatory response

We started to evaluate the role of ErsA in virulence and pathogenicity of *P. aeruginosa* by using an *in vitro* infection system based on pulmonary cell lines. Infections were performed with two prototype laboratory strains, PA14 and PAO1, which are hyper-virulent and moderately virulent, respectively, along with their knock-out counterparts (Δ*ersA*), or with *ersA*-overexpressing PAO1 strains in a wild-type genetic background. The overexpression of ErsA by the vector pGM-*ersA* mimics the increase in ErsA levels induced by the σ^22^-mediated envelope stress response, producing a fivefold-increase of the sRNA levels (29).

We evaluated the influence of ErsA on the cytotoxicity elicited by *P. aeruginosa* during the infection of proliferating CF bronchial epithelial cells IB3-1 through the 3-(4,5-dimethylthiazol-2-yl)-5-(3-carboxymethoxyphenyl)-2-(4-sulfophenyl)-2H-tetrazolium (MTS) colorimetric assay. We exposed pulmonary IB3-1 cells to PA14 or PAO1 wild-type strains and their corresponding Δ*ersA* mutants and followed cell viability by MTS assay during the subsequent 3 hrs. Non-exposed cells were used as control of unaffected viability in the experiment time window (Figs 1A and B). Infection of IB3-1 cells with the PA14 wild-type strain (Fig 1A) caused significant cytotoxic effects, 30% of cell death (loss of viable cells), compared to the non-infected control at the first time point post-infection (60 min). At later time points of 120 and 180 min, the percentage of dead cells was supposed to mirror the pattern of the ratio between death and proliferation rates. In the same conditions, PA14 Δ*ersA* strain showed an interestingly different trend of infection-induced cytotoxicity. Following the killing effects detected at 60 min post-infection, the percentage of dead IB3-1 cells significantly and progressively decreased relatively to PA14 wild-type infection at time points of 120 and 180 min, indicating a cytotoxicity attenuation of PA14 Δ*ersA* strain (Figs 1A and B).

**Fig 1.**
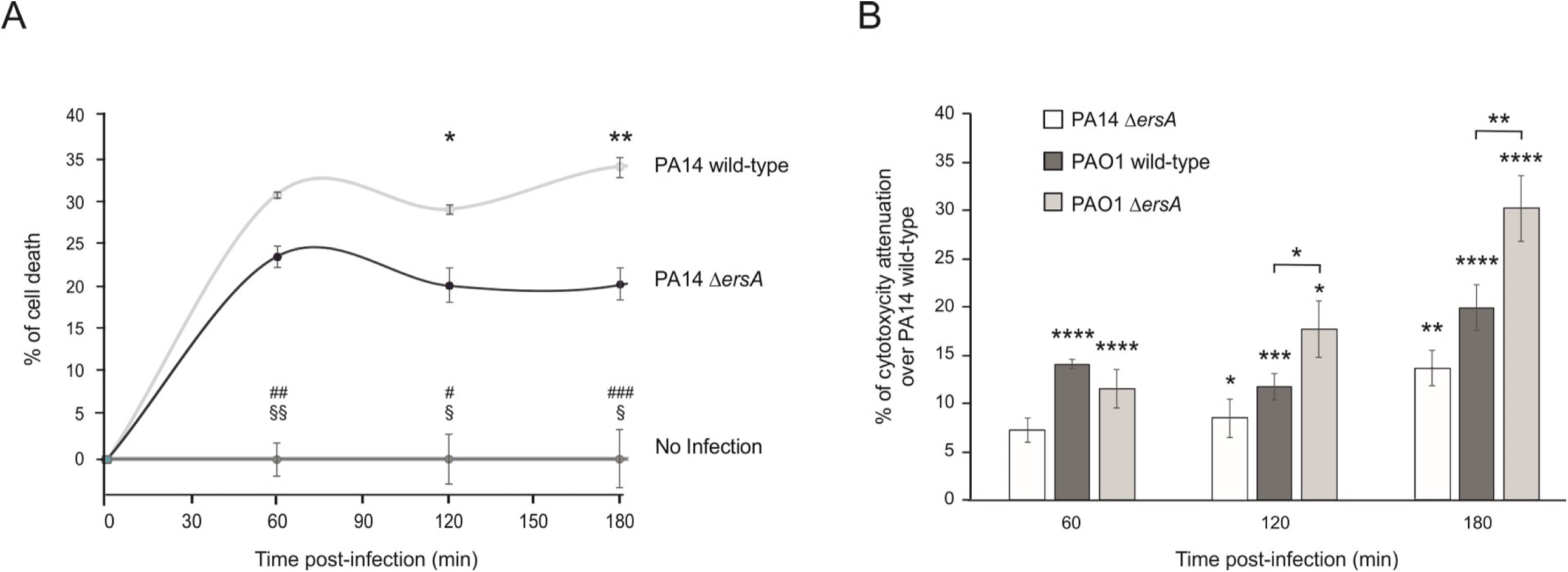
Deletion of ErsA results in decreased *P. aeruginosa*–induced cytotoxicity of pulmonary cells. (A) Time course of cell death of CF bronchial epithelial cells after bacterial infection with *P. aeruginosa* PA14 wild-type and Δ*ersA*. Viability of IB3-1 cells uninfected (No Infection) or infected with a MOI=100 (PA14 wild-type and PA14 Δ*ersA*) was analyzed by MTS assay. At each time point, results are plotted as the ratio of the average values in infected (blank-subtracted) cells to the uninfected cells. The data are pooled from three independent experiments and are represented as mean ± SEM. Significance by one-way *ANOVA* with post-hoc Tukey’s HSD is indicated as follows. PA14 wild-type *vs* Δ*ersA*: *p<0.05 and **p<0.01; PA14 wild-type *vs* No Infection: #p<0.05, ^##^p<0.01 and ^###^p<0.001; PA14 Δ*ersA vs* No Infection: ^§^p<0.05 and ^§§^p<0.01. (B) Relative viability percentage of IB3-1 cell after bacterial infection as measured by MTS assay. Cytotoxicity attenuation (%) of PAO1 wild-type, PAO1 Δ*ersA*, and PA14 Δ*ersA* is shown respect to the PA14 wild-type strain during infection of IB3-1 cells with a MOI=100. At each time point after infection, results are plotted as the ratio of the average values in infected (blank-subtracted) cells to the uninfected cells. At the indicated time points after infection, the ratio of the average values in infected (blank-subtracted) cells to the uninfected cells has been determined. Results are shown as the difference of the ratios between each strain and PA14 wild-type. Data are pooled from three independent experiments and are represented as mean ± SEM. *p<0.05, **p<0.01, ***p<0.001 and ****p<0.0001 in the one-way *ANOVA* with post-hoc Tukey’s HSD. Significance of each strain *vs* PA14 wild-type is indicated above single histograms.

The infection-induced cytotoxicity was assessed for PAO1 wild-type and PAO1 Δ*ersA* in the same experiments described above. Fig 1B reports the results in terms of percentage of cytotoxicity attenuation compared to PA14 wild-type infections shown in Fig 1A, namely the differences, at each time point, between the percentage of dead IB3-1 cells elicited by PA14 wild-type, the most virulent strain of the panel, and those for the other strains. As expected, the PAO1 wild-type strain showed an attenuated phenotype compared to PA14 wild-type. At each time point, the percentages of dead IB3-1 cells were significantly lower than during the infection with PA14 wild-type (i.e. a positive % of attenuation; Fig 1B). The same was true for the PAO1 Δ*ersA* strain but with an extremely relevant difference. Indeed, the percentage of cytotoxicity attenuation resulted significantly higher than PAO1 wild-type at 120 min and further increased at 180 min post-infection. These results for PAO1 are consistent with those presented above for PA14 and strongly suggest that the loss of ErsA affects the cytotoxic potential of *P. aeruginosa*.

We then evaluated the impact of ErsA on the inflammatory response of the IB3-1 cells by monitoring the infection-induced secretion of the pro-inflammatory marker interleukin IL-8, the major chemokine associated with neutrophil extravasation from the vasculature into the lumen of the airways when respiratory epithelial cells are exposed to *P. aeruginosa*. To minimize negative effects that could perturb a robust evaluation of IL-8 secretion, we set out to expose the IB3-1 cells to *P. aeruginosa* strains at a multiplicity of infection (MOI) 10^3^-fold lower than the infection experiments described above. However, PA14-based strains at this MOI still caused relevant cell death, suffering, and detachment from the plastic surface. Such effects were negligible following infection with PAO1-based strains that thus were chosen for the subsequent analyses as follows. IB3-1 cells were exposed to bacteria for 2 hrs, washed, supplemented with amikacin to kill bacteria, and further incubated in the presence of fresh medium with amikacin. Uninfected cells treated and incubated in the same conditions were used as a control of non-stimulated IL-8 production. The amounts of IL-8 released in the supernatants by IB3-1 cells at 24 hrs post-infection were measured through enzyme-linked immunosorbent assay (ELISA). Our results showed that the infection of IB3-1 cells with *P. aeruginosa* PAO1 Δ*ersA* causes a significant decrease in the secretion of IL-8 of about 37% compared to the infection with PAO1 wild-type (Fig 2A). Consistently, IB3-1 cells infected with the ErsA-overexpressing PAO1 strain showed an increase of IL-8 secretion of about 30% compared to those infected with the PAO1 strain harboring the empty vector pGM931 (Fig 2B). Given the difference in the growth conditions of the marker-less strains (PAO1 wild-type and Δ*ersA*, Fig 2A) and the vector-harboring strains (PAO1 pGM931 and pGM-*ersA*, Fig 2B), the absolute amounts of secreted IL-8 are not comparable in these two experimental settings. In the latter case, the presence of carbenicillin for vector maintenance, and arabinose for induction of ErsA expression, may influence the bacterial physiology and, as a consequence, affect the degree of the IB3-1 inflammatory response. In any case, higher ErsA levels in bacterial cells result in a significantly higher secretion of IL-8 in infected IB3-1 cells, as can be concluded from the analysis of both these experimental sets.

**Fig 2.**
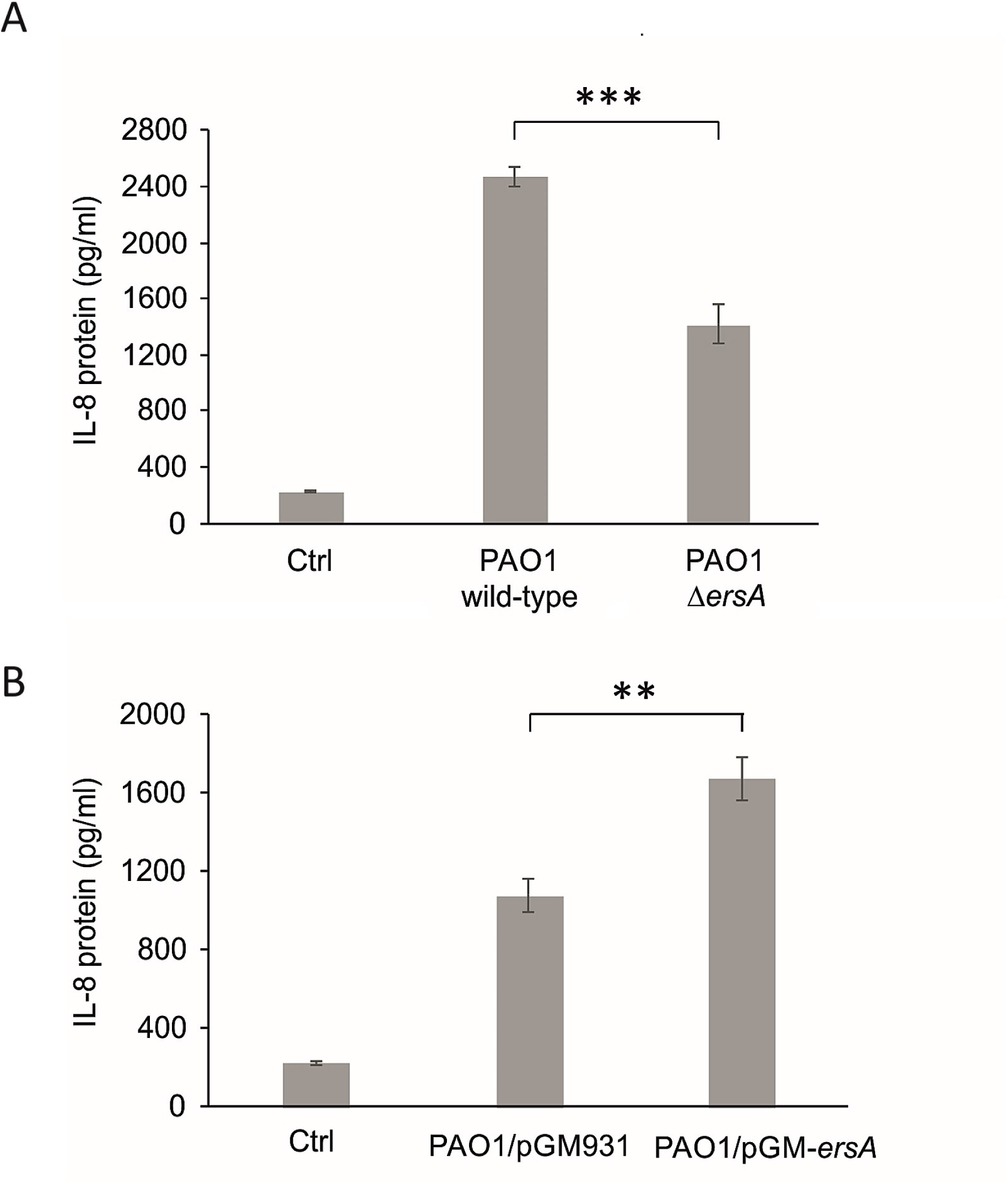
ErsA levels influence the pro-inflammatory response in pulmonary cells. Inflammatory response of CF bronchial epithelial cells after stimulation with (A) *P. aeruginosa* PAO1 wild-type and PAO1 Δ*ersA* deleted mutant strains and (B) *P. aeruginosa* PAO1 strain harboring the empty vector pGM931, or the sRNA overexpressing vectors pGM-*ersA*. IL-8 was evaluated by ELISA in supernatants of IB3-1 cells 24 hrs post-infection (MOI=0.1). Uninfected IB3-1 cells were used as control (Ctrl). Data are represented as mean ± standard error of the mean (SEM). The data are pooled from three independent experiments. *p<0.05, **p<0.01, ***p<0.001 in the Student’s t-test.

Overall, these *in vitro* infection models indicated that ErsA regulatory role impacts *P. aeruginosa*-induced cytotoxicity and contributes to the stimulation of the pro-inflammatory response of infected epithelial cells.

### The deletion of ErsA impairs virulence and decreases pro-inflammatory response in a murine model of airways infection

To further assess the ErsA involvement in the pathogenicity of *P. aeruginosa*, we observed the infection outcomes in two groups of immunocompetent C57BL/6NCrlBR mice whose lungs were inoculated with either PAO1 wild-type or PAO1 Δ*ersA* strain. For this assessment, we followed a protocol of airways infection in which mice are inoculated with bacterial cells embedded in agar beads and monitored for 13 days. We selected this model of infection since it allows the simultaneous analysis of the effects of bacterial mutations on both acute and chronic infection rates. Indeed, agar beads provide microaerobic/anaerobic conditions that allow bacteria to experience a lung environment resembling that of CF (and COPD) patients characterized by thick mucus (13, 38-40). In these conditions, infecting bacteria can either colonize, spread locally and persist in lung establishing a chronic infection, or undertake early systemic dissemination and eventually induce death (acute infection). Alternatively, bacterial cells can be cleared by the host. Depending on virulence, CF airways-adaptation, and the dose of the inoculated bacteria, the fatality rate due to acute infection and the percentage of surviving mice with stable bacterial loads in the lung, signs of chronic infection, can differ considerably in this model of airways infection. For example, the *P. aeruginosa* CF-adapted strain RP73 elicits very low mortality and a high percentage of chronic infection (about 80%) while PAO1 causes significantly higher acute infection-induced mortality and lower chronicity rates (about 15-20 %), thus showing more virulence and lower resilience to host-mediated clearance (38).

Since the PAO1 Δ*ersA* strain showed to be less pro-inflammatory than wild-type in the infection experiments in *vitro* (Fig 2A), in addition to the assessment of bacterial chronic colonization *vs* clearance, surviving mice were also inspected for immune system-activation markers, both in lung and bronchoalveolar lavage fluid (BALF). Specifically, neutrophil and macrophage titers were measured in BALFs while the levels of two pro-inflammatory mediators of the response of airway epithelial cells to *P. aeruginosa* infection, namely the keratinocyte chemoattractant KC (homologous to human IL-8), and the monocyte secretory protein JE (homologous to human monocyte chemoattractant protein-1 MCP-1), were assessed in lung homogenates.

As shown in Fig 3A, the PAO1 Δ*ersA* mutant strain caused significantly lower mortality compared with the PAO1 wild-type counterpart with an infection fatality rate of 0% for Δ*ersA* mutant *vs* 50% for wild-type. Hence, the loss of ErsA resulted in a strong decrease in virulence. Conversely, the incidence of chronic colonization in surviving mice at 13 days post-infection did not differ significantly between Δ*ersA* and wild-type strains (20% *vs* 14% for Δ*ersA* and wild-type strains, respectively; Fig 3B). Likewise, colony-forming unit (CFU) counts were similar in the lung of mice infected with Δ*ersA* mutant and with the wild-type strain (median values of total CFUs: 3.63×10^2^ Δ*ersA* mutant *vs* 3.14×10^2^ wild-type; Fig 3C). The inflammatory response of mice infected by PAO1 Δ*ersA* in terms of leukocytes recruitment in the bronchoalveolar lavage fluid (BALF) was only slightly lower compared to PAO1 wild-type (mean values of total cells: 2.28×10^4^ Δ*ersA* mutant *vs* 2.81×10^4^ wild-type; Fig 3D). However, when chemokines were measured in lung homogenates, we found that PAO1 Δ*ersA* induced significantly lower levels of both KC and JE in comparison to PAO1 wild-type (mean values of KC: 3.21×10^3^ pg/ml Δ*ersA* mutant *vs* 4.07×10^3^ pg/ml wild-type; mean values of JE: 85.7 pg/ml Δ*ersA* mutant *vs* 115.6 pg/ml wild-type; Figs 4A and 4B).

**Fig 3.**
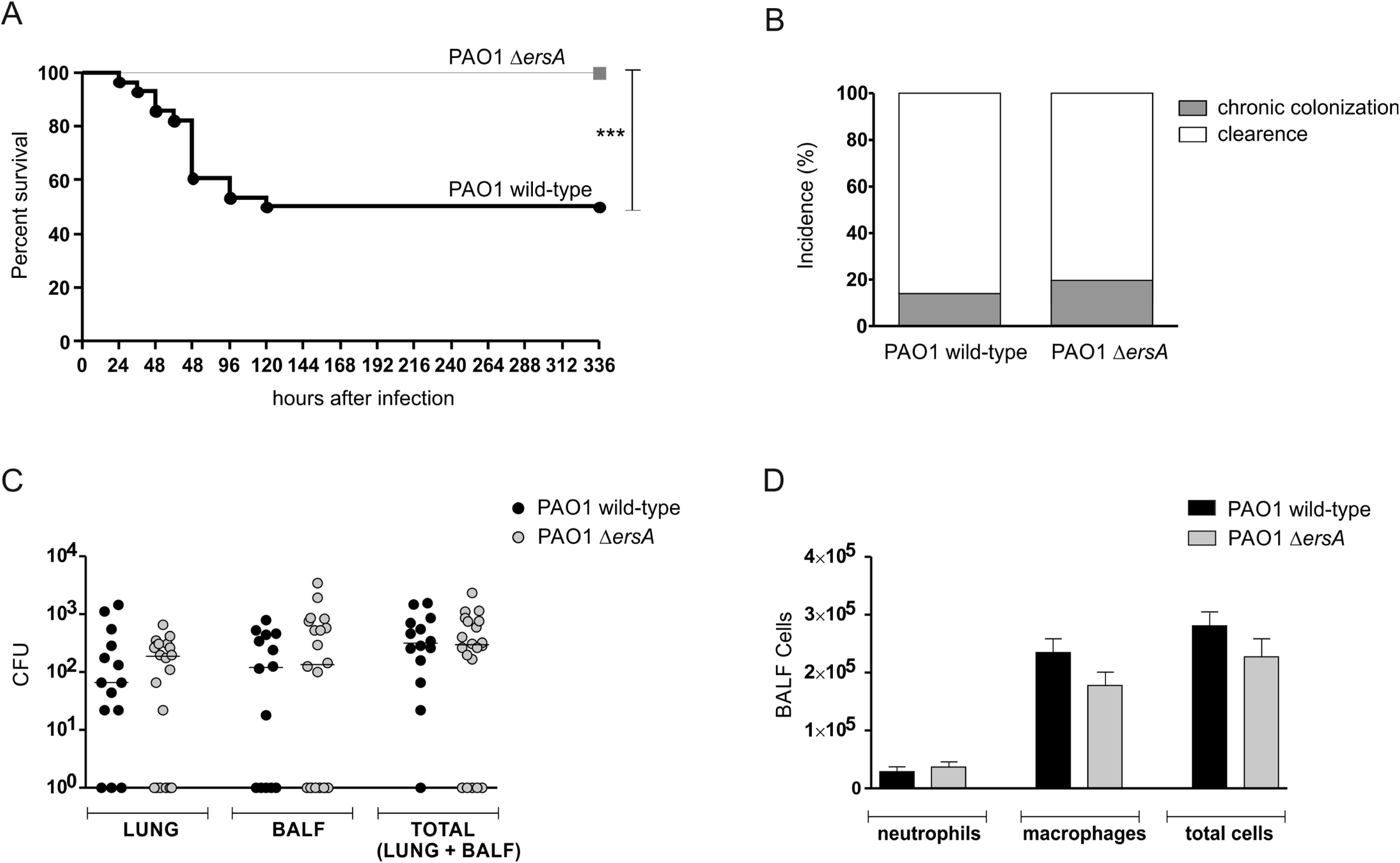
Survival, the incidence of chronic colonization, bacterial burden, and leukocyte recruitment after chronic lung infection by wild-type and Δ*ersA P. aeruginosa* PAO1. C57BL/6NCrlBR mice were infected with 1×10^6^ colony-forming units/lung embedded in agar beads. At day 13 post-infection, mice were sacrificed, bronchoalveolar lavage fluid (BALF) was collected and lungs were excised and homogenized. (A) Survival was evaluated on challenged mice. (B) Clearance (<1000 CFU of *P. aeruginosa* from lung + BALF cultures) and capacity to establish chronic airways infection (≥1000 CFU of *P. aeruginosa* from lung + BALF cultures) were determined on surviving mice. (C) CFUs were evaluated in the lungs and BALF after plating onto tryptic soy agar. Dots represent values of individual mice, and horizontal lines represent median values. (D) Neutrophils, macrophages, and total cells were measured in the BALF. Values represent the mean ± SEM. The data were pooled from at least three independent experiments (n=20-28). ***p<0.001 in the Mantel-Cox test.

**Fig 4.**
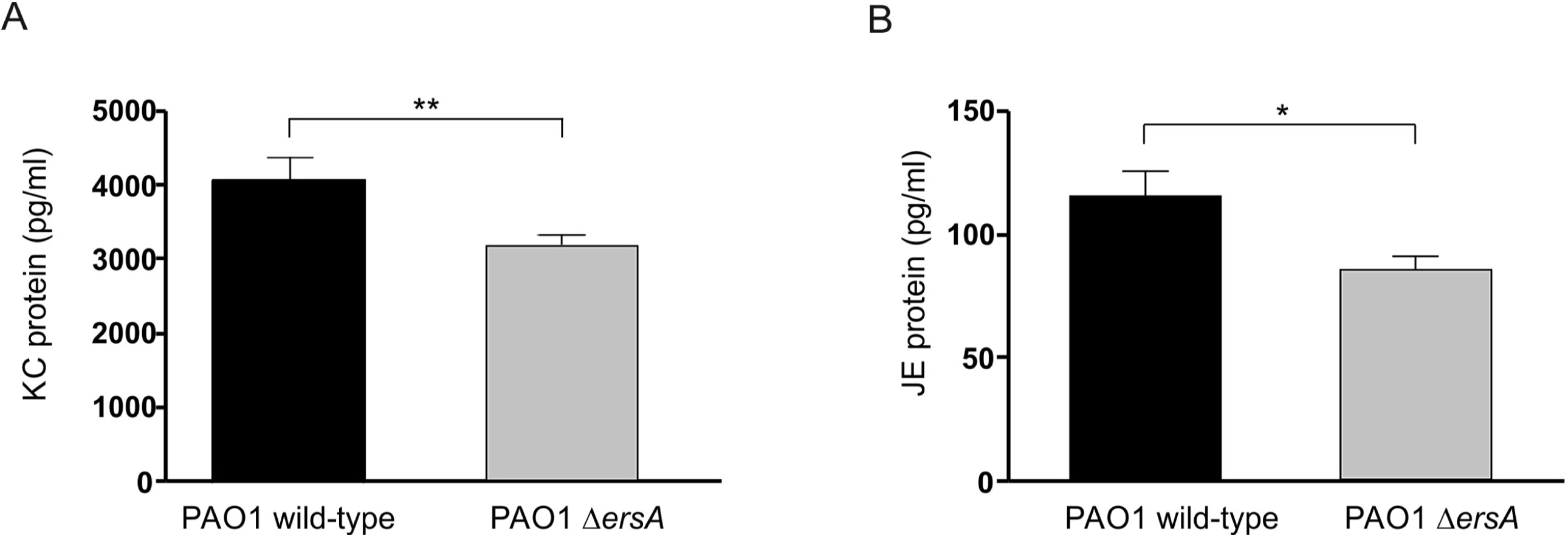
Chemokine levels after lung infection by wild-type and Δ*ersA P. aeruginosa* PAO1. C57BL/6NCrlBR mice were infected with 1×10^6^ colony-forming units/lung embedded in agar beads. At day 13 post-infection, mice were sacrificed, and lungs were excised and homogenized. (A) KC and (B) JE levels were measured by ELISA in the supernatant fluids of lung homogenates. Values represent the mean ± SEM. The data were pooled from at least three independent experiments (n=14-20). *p<0.05, **p<0.01 in the nonparametric two-tailed Mann-Whitney U test.

Overall, these results indicated that the ErsA regulatory function strongly impacts the cascades of events leading to acute infection and lethality and pro-inflammatory response. No influence of ErsA on chronic bacterial colonization of airways is visible in this experimental system.

### Variable ErsA expression in *P. aeruginosa* isolates recovered from human airways chronic infections

Despite the lack of influence on chronic colonization of murine airways, it could be speculated that the attenuation of virulence and a lower stimulation of the immune response potentially deriving from spontaneous deletion, point mutations, or even downregulation of *ersA* gene might favor *P. aeruginosa* persistent lifestyle in human lungs. In PAO1 and PA14 strains, we showed that ErsA expression is strictly dependent on the envelope stress-responsive sigma factor σ^22^ (29). Moreover, ErsA levels can be fine-tuned in response to other environmental cues by additional transcription factors (29). Comparison of the *ersA* gene along with its upstream DNA region in several *P. aeruginosa* isolates from clinical and environmental niches indicated high and extended sequence conservation, including also the -10/-35 core promoter motifs recognized by the RNA polymerase containing σ^22^ (29). This preliminary observation hinted at the possibility that the *ersA* gene itself and its expression responsiveness might be conserved independently of the origin of the *P. aeruginosa* isolates. Nevertheless, it may be feasible that patho-adaptive mutations leading to *ersA* down-regulation contributed to the chronic colonization of the human lung by *P. aeruginosa*.

To address this issue, the presence of *ersA* gene and its expression levels were assessed in a panel of 31 *P. aeruginosa* strains isolated in respiratory samples from CF and COPD patients collected during intermittent or chronic infections at different stages, and compared with those in 5 *P. aeruginosa* isolates from environmental habitats (13, 41, 42) using as references the PAO1 strain. Detection by PCR of the e*rsA* gene and Northern blot analyses are shown in Fig 5 and summarized in S1 Table. The *ersA* gene was detected in 30 out of the 31 clinical isolates and in all environmental strains. In about 55.6% of the analyzed strains, the expression levels of ErsA were not significantly different from those detected in PAO1. This set included the 5 environmental strains and 15 clinical isolates. In another assembly of 13 CF strains, ErsA was significantly down-regulated: in 10 strains from 2 to 7.5 folds, in 2 strains, MI2-3 and TR1, it was strongly downregulated of 28 and 120 folds, respectively, and in one strain, MI1-3, no expression was detected because of the loss of the *ersA* gene as mentioned above. In only three clinical strains ErsA was significantly upregulated from 2 to 3 folds. Hence, these results indicated that, in 13 out of 31 clinical strains analyzed (about 42%), ErsA was moderately to strongly downregulated (in one case lost) relative to the environmental strains that showed to express ErsA at levels comparable to PAO1. This would suggest that ErsA expression is under selective pressure in the CF lung and that mutation(s) resulting in ErsA down-regulation might contribute in some cases to *P. aeruginosa* patho-adaptation during CF chronic infections.

**Fig 5.**
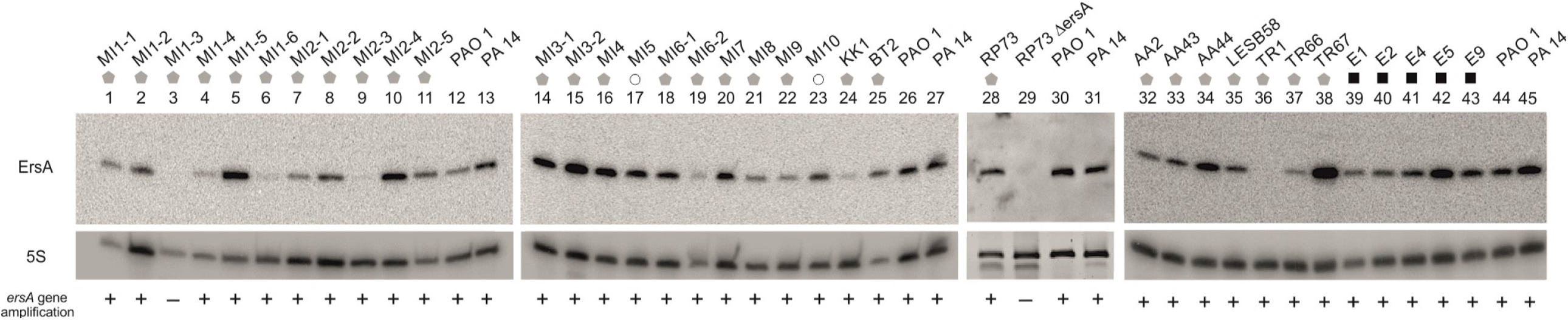
Dissemination of the *ersA* gene and its expression levels in a collection of *P. aeruginosa* isolates. Bacterial strains from CF (gray pentagons), COPD patients (white circles), and environmental isolates (black squares) are indicated on top. After overnight growth at 37°C on BHI-agar plates, culture samples were processed for genomic DNA extraction and total RNA purification and analysis by Northern blot, and for. PAO1 and PA14 were used as control strains. The presence (+) or absence (-) of the *ersA* gene is indicated below each Northern Blot. of Northern blot lane. The relative abundance of ErsA in each isolate was calculated to the reference strain PAO1 after normalization to 5S RNA.

### ErsA can contribute to *P. aeruginosa* adaptation to long-term antibiotic treatment

To further investigate the potential role of ErsA in the *P. aeruginosa* adaption to the CF lung environment, we considered the emergence of antibiotic resistance that is observed frequently in *P. aeruginosa* isolates from CF patients following prolonged antibiotic treatment. To this end, we generated a knock-out *ersA* mutant in RP73, one member of the panel of *P. aeruginosa* clinical isolates that we inspected for ErsA expression (Fig 5, lanes 28 and 29). We selected RP73 because it was isolated from a CF patient 16.9 years after the onset of chronic colonization and showed acquired multi-drug resistance to amikacin, gentamicin, ceftazidime, imipenem, and meropenem (43). The RP73 Δ*ersA* mutant was tested for the Minimum Inhibitory Concentrations (MICs) of seven antibiotics, commonly used in the clinical practice, to which RP73 is resistant. As shown in Table 1, RP73 Δ*ersA* showed to be sensitive to ceftazidime (MIC from 16 in RP73 to 8 µg/ml) and cefepime (MIC from >= 64 in RP73 to 8 µg/ml), and intermediate to meropenem (MIC from >= 16 in RP73 to 4 µg/ml). Furthermore, RP73 Δ*ersA* showed a decrease of MIC from 2 to 1 µg/ml for ciprofloxacin. These results suggest that ErsA could contribute to *P. aeruginosa* adaptation to long-term antibiotic treatment undergone by CF patients.

**Table 1.**
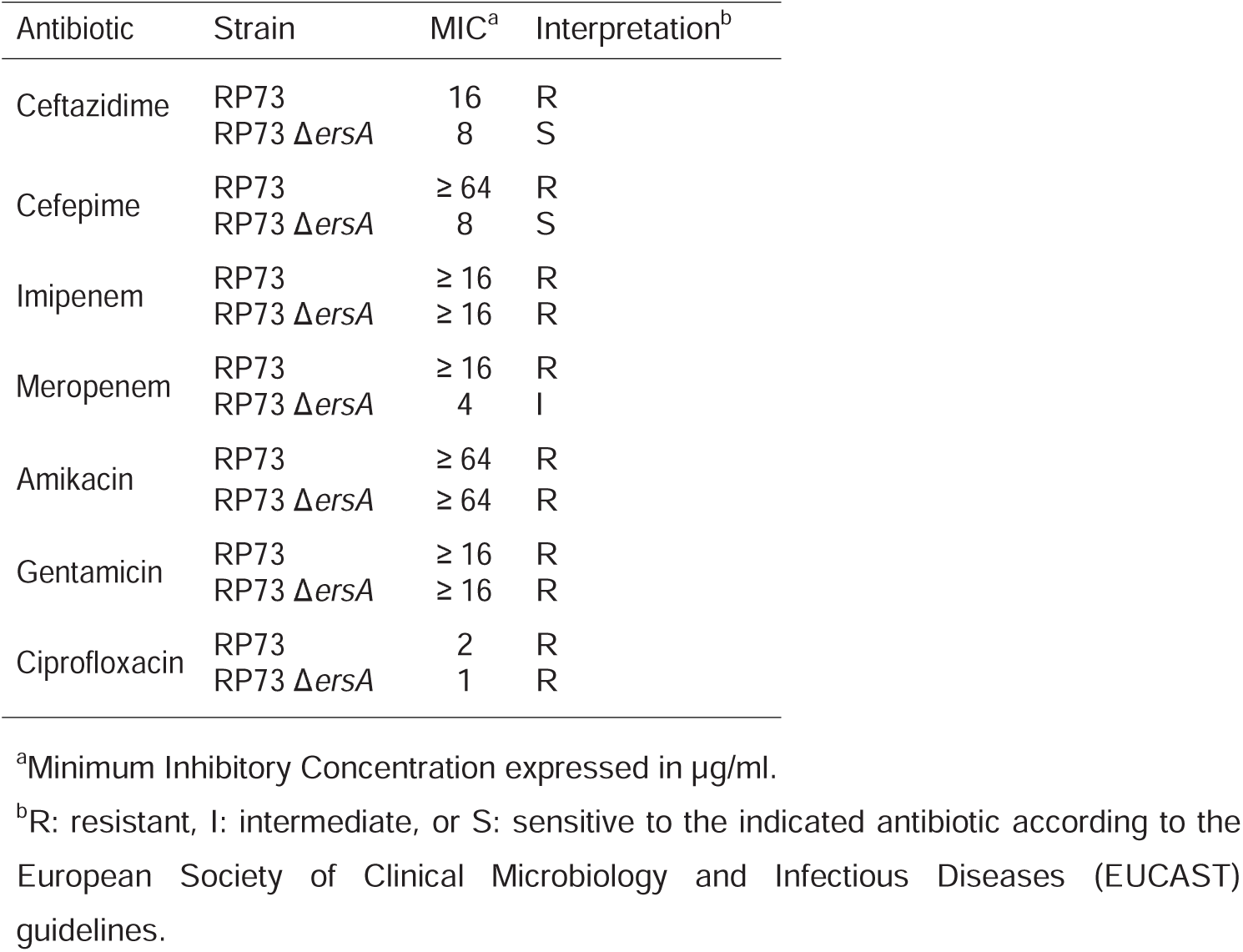
Antibiotic sensitivity of RP73 and RP73 Δ*ersA* strains.

## Discussion

We investigated the regulatory role of ErsA in the pathogenicity and adaptation of *P. aeruginosa* during the infection of the airways. Before this study, several features of ErsA suggested its involvement in the interaction with the host. ErsA regulates EPS production (16) and positively influences biofilm formation and maturation (17). Besides, ErsA responds to cues that are related to airways infection, both at early and late stages, such as a shift from room to body temperature, oxygen availability, iron concentration, and σ^22^-mediated envelope stress response (29), the latter strongly involved in pathogenicity regulation in Gram-negative bacteria (21). ErsA is also involved in the resistance to carbapenem antibiotics through the negative regulation of the porin OprD (37). The results herein presented indicate that the regulatory function exerted by ErsA is relevant in the airways for the progress of *P. aeruginosa* acute infection and might also endure remodeling during the adaptive process leading to *P. aeruginosa* persistence in CF lungs.

The lower *in vitro* cytotoxicity induced by the *P. aeruginosa* Δ*ersA* mutants is consistent with the strong decrease of the fatality rate due to acute infection observed for the PAO1 Δ*ersA* strain. Remarkably, we administered to mice the median lethal dose (LD_50_) of 1×10^6^ CFU for PAO1, which was completely ineffective in the case of PAO1 Δ*ersA*. Besides, there is another consistency of results between the *in vitro* and *in vivo* infection experiments: the loss of ErsA determines a lower activation of the innate immune response, measured in terms of levels of the NF-kB-dependent pro-inflammatory mediators IL-8 (*in vitro*), and KC and JE (*in vivo*). Hence, the regulatory role of ErsA impacts both the virulence of the acute infection and the innate immune response. To explain these results, we first speculated that ErsA could positively regulate in response to lung environment invasive functions that can also act as PAMPs (e.g. flagella, LPS, T3SS, and ExoS) (10), or the expression of non-PAMP invasive functions and PAMPs products. Alternatively, the significant impairment of PAO1 Δ*ersA* in biofilm formation and maturation (30) could justify the simultaneous involvement of ErsA in acute infection and immune response activation. Previous relevant studies (5, 6, 9) have evidenced that, at initial stages of *P. aeruginosa* acute infection, biofilm aggregates with a canonical matrix composed of Psl, Pel, alginate, and eDNA (4) assemble on airways mucosal surface and trigger both a dramatic remodeling of the apical membrane (i.e. protrusions) of epithelial cells and NF-kB-dependent activation of the innate immune response. Neither protrusion formation nor NF-kB activation was observed upon binding of individual bacteria to epithelial cells. This strongly indicated that biofilm formation is a key *P. aeruginosa* function to initiate acute infection and, through the induced changes in epithelial cell polarity, a danger signal for host cells that warns of an incoming threat (7). On the bases of this model, we suggest that acute virulence attenuation and decreased pro-inflammatory stimulation of PAO1 Δ*ersA* could be due to its defect in biofilm formation and maturation (30). This phenotype induced by the loss of ErsA was attributed to the dysregulation of the expression of AlgC (29, 33) and AmrZ (44, 45), two proteins that play important roles in the post-transcriptional and transcriptional regulation, respectively, of the production of Psl, Pel, and alginate. Furthermore, transcriptomics analysis indicated that the *pelCDEFG* genes for Pel biosynthesis (46), the *ppyR* gene for an activator of the Psl operon coding for the Psl biosynthetic pathway (47), and the *algD* gene for alginate biosynthesis are significantly down-regulated in the PAO1 Δ*ersA* mutant (30).

Specifically, Psl and Pel (referred to as aggregative EPS) are important for initiating and maintaining cell-cell interaction in biofilms, while alginate (referred to as capsular EPS) is instrumental in biofilm maturation, structural stability and, protection from antibiotics (48). To further assess the role of ErsA in the regulation of Psl- and Pel-linked aggregation and adherence, we performed the experiments shown in S1 Fig. As a result of ErsA deletion, aggregation, and adherence potentials of PAO1 are strongly reduced, while they are enhanced when ErsA is overexpressed. Overall, ErsA could participate in the regulation of biofilm formation at the early stages of acute infection. However, this scenario might be wider since i) transcriptomics data (30) suggested that ErsA deletion can affect other aspects of *P. aeruginosa* interaction with its host and ii) at the post-transcriptional level, ErsA could influence the expression of virulence-associated genes other than *algC* and *amrZ*.

The deletion of ErsA did not influence the chronic infection rate of PAO1 in the mouse model. However, virulence attenuation and lower recognition by the immune system showed by PAO1 Δ*ersA* are favorable traits for *P. aeruginosa* chronic infection of CF airways (4, 10). Therefore, we evaluated whether ErsA expression could be downregulated or even deleted in a panel of CF clinical isolates.

A significant proportion (about 42%) of the clinical strains analyzed showed that ErsA was moderately to strongly downregulated (in one case lost) relative to both environmental strains and PAO1. This suggested that the *ersA* gene could be under selective pressure for lowering its expression in a CF context. This phenomenon might contribute in some cases to *P. aeruginosa* patho-adaptation towards low virulence and evasion of the immune system during CF chronic infections (10). We speculate that this putative evolution of ErsA expression occurs in the frame of the remodeling process involving the infection regulatory network, in which σ^22^ is one main component that leads to *P. aeruginosa* adaptation to CF lung (17).

Finally, we found that loss of ErsA induces sensitization to ceftazidime, cefepime, and meropenem in the multidrug-resistant clinical isolate RP73, which was in the group of CF clinical isolates analyzed for ErsA expression. It is worth noting that RP73 was demonstrated to establish long-term infection replacing an initial isolate (RP1), and adapting within CF airways compared to its clonal ancestor RP45 (43). The adaptive microevolution has led RP73 to differentiate significantly from RP45 in terms of virulence and antibiotic resistance, being RP45 more virulent than RP73 and sensitive to amikacin, ceftazidime, imipenem, and meropenem (43). The multi-antibiotic resistance that RP73 has acquired compared to its clonal ancestor RP45, and that is lost in the RP73 Δ*ersA*, reveals an interesting link between ErsA and mechanisms of adaptation to host environment during *P. aeruginosa* chronic infection of CF patients. The emergence and rapid dissemination of antibiotic resistance demand the development of new antibiotics and antivirulence agents (49). These latter compounds directly target virulence factors or virulence regulators. The contribution to acute infection regulation and the acquirement of antibiotic resistance suggest that ErsA may be a candidate target for the development of novel anti-virulence and co-antibiotic drugs.

## Materials and methods

### Ethics Statement

The study on human *P. aeruginosa* isolates from Hannover has been approved by the Ethics Commission of Hannover Medical School, Germany (41). The patients and parents gave oral informed consent before the sample collection. Approval for storing the biological materials was obtained by the Ethics Commission of Hannover Medical School, Germany. The study on human *P. aeruginosa* isolates from the Regional CF Center of Lombardia was approved by the Ethical Committees of San Raffaele Scientific Institute and Fondazione IRCCS Ca’ Granda, Ospedale Maggiore Policlinico, Milan, Italy, and written informed consent was obtained from patients enrolled or their parents according to the Ethical Committees rules, under the laws of the Italian Ministero della Salute (approval #1874/12 and 1084/14) (42).

Animal studies strictly followed the Italian Ministry of Health guidelines for the use and care of experimental animals. This study was performed following protocols approved by the Institutional Animal Care and Use Committee (IACUC, protocol #789) of the San Raffaele Scientific Institute (Milan, Italy).

### Bacterial strains and culture conditions

*P. aeruginosa* strains PAO1 (26), PA14 (27) and RP73 (41, 43), and the corresponding deleted mutants PAO1 Δ*ersA*, PA14 Δ*ersA* (29) and RP73 Δ*ersA* were grown at 37 °C in Luria-Bertani rich medium at 120 rpm. sRNA-overexpressing strains PAO1/pGM-ersA and PA14/pGM-ersA and their empty vector-harboring control strains PAO1/pGM931 and PA14/pGM931 (29) were grown with the addition of 300 μg/ml carbenicillin. For *P*_*BAD*_ induction in vector plasmid pGM931, arabinose was added to a final concentration of 10 mM. The RP73 Δ*ersA* mutant strain was generated from the MDR-RP73 isolate using a method of marker-less gene replacement (50) improved for *P. aeruginosa* using oligos and molecular techniques as described previously (29), and cloning in the tetracycline-resistant harboring plasmid pSEVA512S to allow selection of exconjugant on 30 μg/ml tetracycline.

### Bacterial isolates analysis

Bacterial isolates (13, 41, 42) were plated on 1.5% Brain Heart Infusion (BHI)-agar plates and grown overnight at 37°C. Culture samples were taken and processed for genomic DNA and total RNA extraction as described previously (28). PAO1 and PA14 strains treated in the same conditions were used as controls.

Oligos CGAATGGCTTGAGCCCTTCGATGCT/AAAAAAAACCCCGAGCTTCGTA and TGTCGTCAGCTCGTGTCGTGA/ATCCCCACCTTCCTCCGGT were used for PCR-amplification of the genomic region containing the *ersA* and 16S (as positive PCR-control) *loci*, respectively. Northern blot analyses were performed as described previously (28). Briefly, DNA oligonucleotide probes were 5’ end-labeled with [γ-32P]ATP (PerkinElmer, NEG502A) and T4 polynucleotide kinase (Promega, M4103) according to the manufacturer’s instruction. Oligo CCCGAGCTTCGTATGGGG and GGAGACCCCACACTACCATCGGCGATG were used to probe ErsA and 5S RNA, respectively. Radioactive bands were acquired after exposure to phosphor screens using a Typhoon(tm) 8600 variable mode Imager scanner (GE Healthcare BioSciences) and visualized with image-Quant software (Molecular Dynamics). The intensities of the bands were quantified using Li-cor Image Studio Lite. The signal of ErsA was normalized to those of 5S RNA in the same lane. For each clinical isolate, the relative abundance of ErsA was calculated comparing to the reference strain PAO1.

### Cytotoxicity and IL-8 secretion in human CF respiratory cells

IB3-1 cells, an adeno-associated virus-transformed human bronchial epithelial cell line derived from a CF patient (ΔF508/W1282X) and obtained from LGC Promochem, were grown as described previously (51). Cell viability was evaluated using the MTS-based CellTiter 96® AQ_ueous_ One Solution Cell Proliferation Assay kit (Promega, G3582), which determines viable cell number measuring the conversion at 490 nm of 3-(4,5-dimethylthiazol-2-yl)-5-(3-carboxymethoxyphenyl)-2-(4-sulfophenyl)-2H-tetrazolium (MTS) to formazan by the dehydrogenase enzyme of the intact mitochondria of living cells. In a 96-well plate, triplicates of IB3-1 cells were infected with *P. aeruginosa* strains at a multiplicity of infection (MOI) of 100, in a final volume of 100 µl. Immediately after infection, 20 µl of the CellTiter 96® AQ_ueous_ One Solution Reagent were added directly to culture and control wells. MTS was also added to non-infected cells and in wells containing only the same bacterial load in absence of IB3-1 cells (blank/control). According to the manufacturer’s instructions, plates were incubated at 37°C with 5% CO_2,_ read at 490 nm at different time-points, and returned to the incubator for further color development. The average measurement of infected cells was subtracted from the average of the corresponding blank. The relative percentages of cell death or cell viability were calculated as the ratio between the average value in normalized infected cells (blank-subtracted) and the uninfected cells. The stimulation of the host inflammatory response was evaluated by monitoring the secretion of the pro-inflammatory interleukin IL-8 as described previously (51). Briefly, after infection with *P. aeruginosa* strains at an MOI of 0.1, IB3-1 cells were incubated at 37°C with 5% CO_2_ for 2 hrs, washed with PBS supplemented with 1 mg/ml amikacin, and incubated in presence of fresh medium supplemented with 1 mg/ml amikacin. Uninfected cells treated and incubated in the same conditions were used as a control of non-stimulated IL-8 production. Released IL-8 was determined in supernatants collected at 24 hrs using an ELISA kit (Biosource Europe and R&D Systems), according to manufacturer’s instructions.

### Agar-beads preparation

The agar-beads mouse model was used (39, 40). An aliquot of wild-type or Δ*ersA P. aeruginosa* PAO1 strains from glycerol stocks was streaked for isolation on trypticase soy agar (TSA) and incubated at 37°C overnight. One colony was picked from the plate and used to inoculate 5 ml of tryptic soy broth (TSB) and placed in a shaking incubator at 37°C 200 rpm overnight. The overnight bacterial suspension was diluted to 0.15 OD/ml in 20 ml of TSB / flask and grown for 4 hrs at 37°C at 200 rpm, to reach the log phase. The bacteria were pelleted by centrifugation (2,700 g, 15 min, 4°C) and resuspended in 1 ml PBS (pH 7.4). A starting amount of 2×10^9^ CFUs of *P. aeruginosa* was used for inclusion in the agar-beads prepared according to the previously described method (39, 40, 52). Bacteria were added to 9 ml of 1.5% TSA (w/v), prewarmed to 50°C. This mixture was pipetted forcefully into 150 ml heavy mineral oil at 50°C and stirred rapidly with a magnetic stirring bar for 6 min at room temperature, followed by cooling at 4°C with continuous slowly stirring for 35 min. The oil-agar mixture was centrifuged at 2,700 g for 15 min to sediment the beads and washed six times in PBS. The size of the beads was verified microscopically and only those preparations containing beads of 100 μm to 200 μm in diameter were used as inoculum for animal experiments. The number of *P. aeruginosa* CFUs in the beads was determined by plating serial dilutions of the homogenized bacteria-bead suspension on TSA plates. The inoculum was prepared by diluting the bead suspension with PBS to 2×10^7^ CFUs/ml, to inoculate about 1×10^6^ CFU/50μl. *P. aeruginosa* beads were prepared the day before inoculation, stored overnight at 4°C for a maximum of two days. The number of *P. aeruginosa* CFUs in the beads inoculated was determined by plating serial dilutions of the homogenized bacteria-bead suspension on the day of the infection.

### Mouse model of chronic *P. aeruginosa* lung infection

Immunocompetent C57BL/6NCrlBR male mice (8-10 weeks of age) were purchased from Charles River (Calco, Italy), shipped in protective, filtered containers, transported in climate-controlled trucks, and allowed to acclimatize for at least two days in the stabulary before use. Mice were maintained in the biosafety level 3 (BSL3) facility at San Raffaele Scientific Institute (Milano, Italia) where three-five mice per cage were housed. Mice were maintained in sterile ventilated cages. Mice were fed with standard rodent autoclaved chow (VRFI, Special Diets Services, UK) and autoclaved tap water. Fluorescent lights were cycled 12h on, 12h off, and ambient temperature (23±1°C) and relative humidity (40-60%) were regulated.

For infection experiments, mice were anesthetized by an intraperitoneal injection of a solution of Avertin (2,2,2-tribromethanol, 97%) in 0.9% NaCl and administered at a volume of 0.015 ml/g body weight. Mice were placed in a supine position. The trachea was directly visualized by ventral midline, exposed and intubated with a sterile, flexible 22-g cannula attached to a 1 ml syringe. An inoculum of 50 μl of agar-bead suspension was implanted via the cannula into the lung. After inoculation, all incisions were closed by suture.

Infections and sacrifices were all performed in the late morning. Besides, in all the experiments, mice had been subdivided according to the bodyweight to have similar mean in all the groups of treatment.

Mice were monitored daily for coat quality, posture, attitude, ambulation, hydration status, and bodyweight. Mice that lost >20% bodyweight and had evidence of severe clinical diseases, such as scruffy coat, inactivity, loss of appetite, poor locomotion, or painful posture, were sacrificed before the termination of the experiments with an overdose of carbon dioxide. Gross lung pathology was checked. After 13 days post-infection, bronchoalveolar lavage fluid (BALF) was collected and lungs were aseptically excised.

BALF was extracted with a 22-gauge venous catheter, ligated to the trachea to prevent backflow. The lungs were washed with three one ml of RPMI-1640 (Euroclone) with protease inhibitors (Complete tablets, Roche Diagnostic) and pooled. Quantitative bacteriology on BALF was performed by plating serial dilution on TSA. Total cells present in the BALF were counted using an inverted light optical microscope after diluting an aliquot of the BALF 1:2 with Tuerk solution in a disposable counting chamber. BALF cells were centrifuged at 330 g for 8 min at 4°C. If the pellet was red, erythrocytes were lysed by resuspending the pellet in 250-300 µl of RBC lysis buffer diluted 1:10 in ultra-pure distilled water for 3 min. Then, 2-3 ml PBS were added and cells were centrifuged at 330 g for 8 min at 4°C. The pellet was resuspended in RPMI 1640 10% fetal bovine serum (FBS) at a concentration of 1×10^6^ cells/ml, and an aliquot of 170 µl was pipetted into the appropriate wells of the cytospin and centrifuged at 300 g for 5 min medium brake. Slides were then stained by Diff-Quik staining using a commercial kit (Medion Diagnostics, code: 726443), according to the manufacturer’s instructions. A differential cell count was performed at an inverted light optical microscope.

Lungs were excised aseptically and homogenized in 2 ml PBS added with protease inhibitors using the homogenizer gentleMACSTM Octo Dissociator. One-hundred μl of the homogenates and 10-fold serial dilutions were spotted onto TSA. CFUs were determined after overnight growth at 37°C. Recovery of ≥1000 CFU of *P. aeruginosa* from lung + BALF cultures was considered as evidence of chronic infection.

### Quantification of murine chemokines

Lung homogenates were centrifuged at 16,000 g for 30 min at 4°C, then supernatants were collected and stored at −80°C. Murine KC and JE concentrations were determined in the lung homogenate supernatants by DuoSet® ELISA Development Systems (R&D Systems), according to manufacturer instructions.

### MIC measurement

MICs of antibiotics were determined according to CLSI guidelines (53), as previously described (54). The medium used for the MIC testing was cation-adjusted Mueller-Hinton broth (MH-II broth). MIC testing was run in sterile 96-well Microtiter plates (polystyrene V shape) and analyzed after 20 hrs.

### Statistics

Statistical analyses were performed with GraphPad Prism. Survival curves and incidences of chronic colonization were compared using the Mantel-Cox test and Fisher exact test, respectively. Levels of chemokines, leukocytes, and CFUs were compared using a nonparametric two-tailed Mann-Whitney U test. *P* < 0.05 was considered significant. Significance of the differences in the levels of secreted IL-8 and converted MTT by infected IB3-1 cells were determined using the Student t-test. *P* < 0.05 was considered significant.

## Acknowledgments

The authors thank Prof. B. Tümmler (Hannover Medical School, Hannover, Germany), and Dr. L. Cariani (Cystic Fibrosis Microbiology Laboratory, Fondazione IRCCS Ca’ Granda, Ospedale Maggiore Policlinico, Milano, Italy) for supplying the *P. aeruginosa* strains from CF and COPD patients. S.F. was the recipient of a Postdoctoral Fellowship of the Università degli Studi di Milano.

## Supporting information

**S1 Fig. ErsA influences aggregation and adherence of the *P. aeruginosa* PAO1 strain**. *P. aeruginosa* PAO1 strains with wild-type, deleted, or overexpressed ErsA grown in liquid T-Broth in presence of 40 µg/ml of Congo red. To observe aggregation, strains were inoculated at OD_600_ of 0.05 using glass culture tubes with 10 ml of medium and incubated overnight at 37°C in a roller drum. Carbenicillin and arabinose were added for ectopic expression of ErsA from pGM-*ersA* and the growth of the control culture harboring the pGM931 empty vector. Adherent biomass is noticeable on the culture tube of PAO1 wild-type strain. Conversely, the absence of adherence is shown on the Δ*ersA* strain tube. More abundant aggregation and biofilm are present when ErsA is over-expressed by pGM-*ersA* from wild-type background, compared to the empty vector-harboring control strain PAO1/pGM931 grown in the same conditions.

**S1 Table. Analysis of ErsA expression in a panel of clinical and environmental strains of *P. aeruginosa.***

