## Supplemental Table 1 for "The small RNA ErsA plays a role in the regulatory network of *Pseudomonas aeruginosa* pathogenicity in airways infection"

**S1 Table. Analysis of ErsA expression in a panel of clinical and environmental strains of *P. aeruginosa.***

| Strain | Origin [a] | Reference | Lane [b] | *ersA gene* detection by PCR | Differential *ersA* expression relative to PAO1 | % [c] |
|---|---|---|---|---|---|---|
| MI1-5 | CF | (1) | 5 | + | | |
| MI3-2 | CF | (1) | 15 | + | upregulation | 8.3 |
| TR67 | CF | (2) | 38 | + | | |
| MI1-1 | CF | (1) | 1 | + | | |
| MI1-2 | CF | (1) | 2 | + | | |
| MI2-2 | CF | (1) | 8 | + | | |
| MI2-4 | CF | (1) | 10 | + | | |
| MI2-5 | CF | (1) | 11 | + | | |
| MI3-1 | CF | (1) | 14 | + | | |
| MI4 | CF | (1) | 16 | + | | |
| MI5 | COPD | (1) | 17 | + | | |
| MI6-1 | CF | (1) | 18 | + | | |
| MI7 | CF | (1) | 20 | + | not significantly different | 55.6 |
| MI10 | COPD | (1) | 23 | + | | |
| BT2 | CF | (2) | 25 | + | | |
| RP73 | CF | (3) | 28 | + | | |
| AA43 | CF | (2) | 33 | + | | |
| AA44 | CF | (2) | 34 | + | | |
| E1 | E | (2) | 39 | + | | |
| E2 | E | (2) | 40 | + | | |
| E4 | E | (2) | 41 | + | | |
| E5 | E | (2) | 42 | + | | |
| E9 | E | (2) | 43 | + | | |
| MI1-3 | CF | (1) | 3 | - | | |
| MI1-4 | CF | (1) | 4 | + | | |
| MI1-6 | CF | (1) | 6 | + | | |
| MI2-1 | CF | (1) | 7 | + | | |
| MI2-3 | CF | (1) | 9 | + | | |
| MI6-2 | CF | (1) | 19 | + | | |
| MI8 | CF | (1) | 21 | + | downregulation or no expression | 36.1 |
| MI9 | CF | (1) | 22 | + | | |
| KK1 | CF | (2) | 24 | + | | |
| AA2 | CF | (2) | 32 | + | | |
| LESB58 | CF | (2) | 35 | + | | |
| TR1 | CF | (2) | 36 | + | | |
| TR66 | CF | (2) | 37 | + | | |

[a] CF: Cystic Fibrosis patient; COPD: Chronic Obstructive Pulmonary Disease patient; E: Environment.

[b] Lane number in Figure 5.

[c] Percentage of strains with the indicated ersA expression.
